# Spatially Resolved Metabolites in Stable and Vulnerable Human Atherosclerotic Plaques Identified by Mass Spectrometry Imaging

**DOI:** 10.1101/2022.05.30.494038

**Authors:** Erin H. Seeley, Zhipeng Liu, Shuai Yuan, Chad Stroope, Elizabeth Cockerham, Nabil Rashdan, Alexandra C Finney, Dhananjay Kumar, Sandeep Das, Babak Razani, Wanqing Liu, James Traylor, Oren Rom, Christopher Pattillo, Arif Yurdagul

## Abstract

Impairments in carbohydrate, lipid, and amino acid metabolism drive features of plaque instability. However, where these impairments occur within the atheroma remains largely unknown. Therefore, we sought to characterize the spatial distribution of metabolites within stable and unstable atherosclerosis in both the fibrous cap and necrotic core. Atherosclerotic tissue specimens were scored based on the Stary classification scale and subdivided into stable and unstable atheromas. After performing mass spectrometry imaging (MSI) on these samples, we identified over 850 metabolite-related peaks. Using MetaboScape, METASPACE, and HMDB, we confidently annotated 170 of these metabolites and found over 60 of these were different between stable and unstable atheroma. We then integrated these results with an RNA-sequencing dataset comparing stable and unstable human atherosclerosis. Upon integrating our MSI results with the RNA-seq dataset, we discovered that pathways related to lipid metabolism and long-chain fatty acids were enriched in stable plaques, whereas reactive oxygen species, aromatic amino acid, and tryptophan metabolism were increased in unstable plaques. Acylcarnitines and acylglycines were increased in stable plaques whereas tryptophan metabolites were enriched in unstable plaques. Evaluating spatial differences in stable plaques revealed lactic acid in the necrotic core, whereas pyruvic acid was elevated in the fibrous cap. In unstable plaques, 5-hydroxyindole-acetic acid was enriched in the fibrous cap. Our work herein represents the first step to defining an atlas of metabolic pathways involved in plaque destabilization in human atherosclerosis. We anticipate this will be a valuable resource and open new avenues of research in cardiovascular disease.

**GRAPHICAL ABSTRACT:** 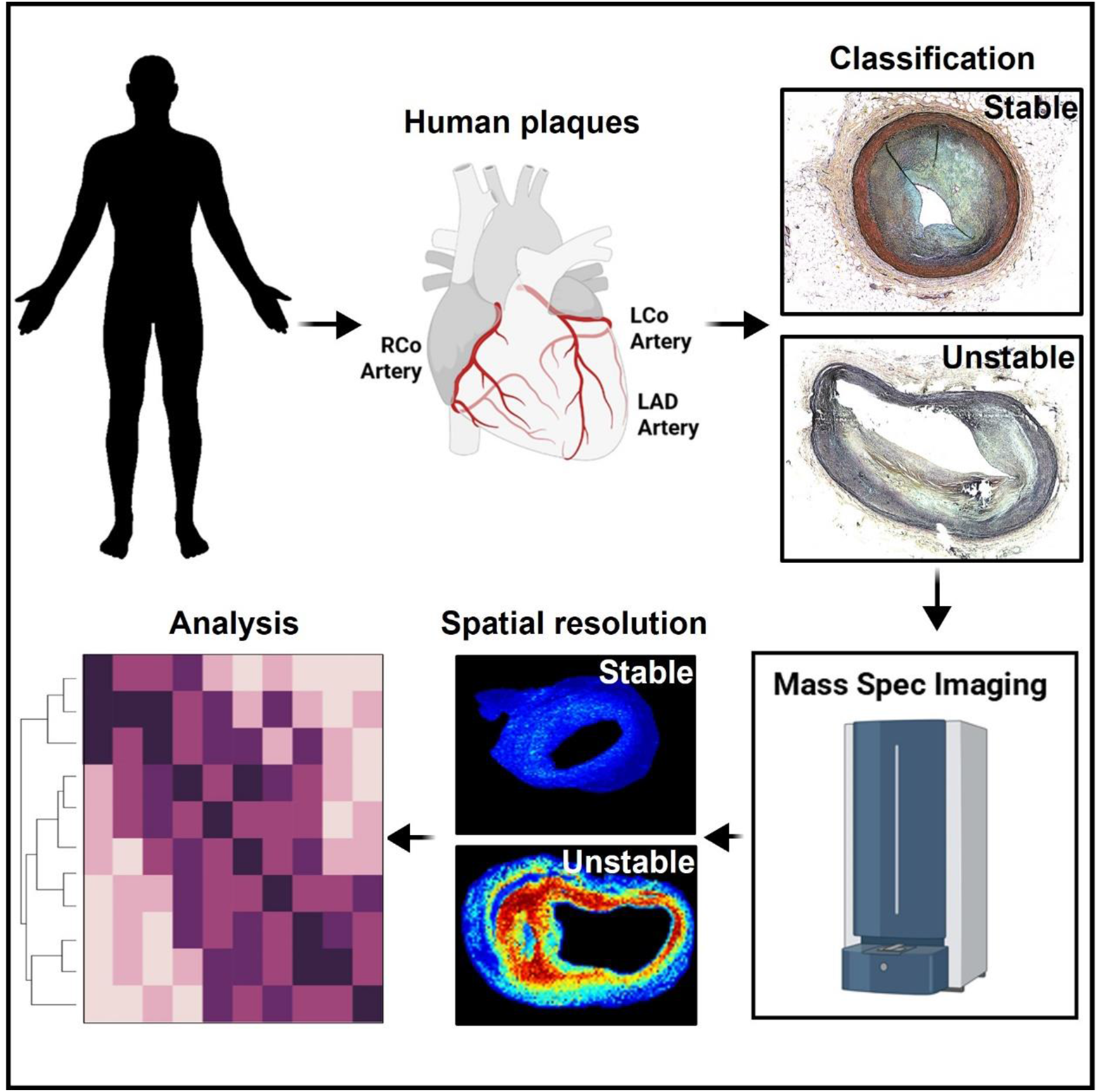

## INTRODUCTION

Despite the considerable advancements in diagnostic techniques and therapeutic strategies, atherosclerotic cardiovascular disease (CVD) remains the leading cause of morbidity and mortality worldwide^1^. Acute thrombogenic events that lead to myocardial infarction and stroke are frequently caused by the rupture of vulnerable plaques, characterized by the presence of a large necrotic core covered by a thin fibrous cap^2,3^. In contrast, stable plaques show thicker collagen-rich fibrous caps with smaller necrotic cores. Understanding these features of stable and vulnerable plaques has provided a framework for much of the work regarding the complexity of mechanisms that eventually lead to CVD-related events.

There is a growing interest in elucidating the mechanisms that govern plaque instability, which has so far highlighted the role of dysregulations in lipid, carbohydrate, and amino acid metabolism^4-9^. Impairments in metabolism play a critical role in all stages of atherosclerosis, from its inception as a benign fatty streak to the precipitation of acute clinical events. However, where these impairments occur in the atherosclerotic plaque has yet to be revealed. This is critically important because lipid deposition, cell composition, and matrix stiffness are heterogeneous throughout human atheromas, particularly in regions of the fibrous cap and near the necrotic core. Furthermore, lesional cells are constantly bombarded with various intrinsic and extrinsic metabolic insults that influence cell phenotype^4,5^. While our understandings of metabolism in atherosclerosis are being elucidated, technical limitations have prevented investigators from determining where these regional impairments in metabolism occur in the plaque.

Mass spectrometry imaging (MSI) is a powerful technology for visualizing the spatial localization and relative abundance of hundreds to thousands of analytes in thin tissue sections without *a priori* knowledge of analytes present in the samples^10^. Imaging is performed using matrix-assisted laser desorption/ionization (MALDI) to interrogate small areas, or pixels, from a tissue surface after adding a chemical matrix to the section, which serves to extract, co-crystallize, and ionize molecules from the sample^11^. With the capability to image thousands of molecules without mass labeling, the information gained from mass spectrometry and the visualization of its spatial distributions makes this platform uniquely suited to characterize the spatial distribution of metabolites in an atherosclerotic plaque. In the present study, we performed MALDI MSI of late-stage, stable and vulnerable human atherosclerotic plaques to visualize the distribution of metabolites in the fibrous cap and necrotic core.

## MATERIALS AND METHODS

The data that support the findings of this study are available from the corresponding authors upon reasonable request. An expanded materials and methods section can be found in the online supplemental file.

### Human atheroma specimen collection and scoring

Human tissue was deemed non-human research by the local IRB due to the exclusive use of postmortem samples. Human tissue was excised postmortem during autopsy and fixed in 4% formaldehyde, embedded in paraffin, then cut into 5 μm sections. Based on stains described in Figure 1, sections were scored into type I (lesions with macrophage foam cells), type II (fatty streaks with intracellular lipid accumulation), type III (increased presence of extracellular lipid pools), type IV (presence of necrotic core and fibrous cap), or type V (similar to type IV except with the presence of calcific nodules or sheets)^12^.

**Figure 1.**
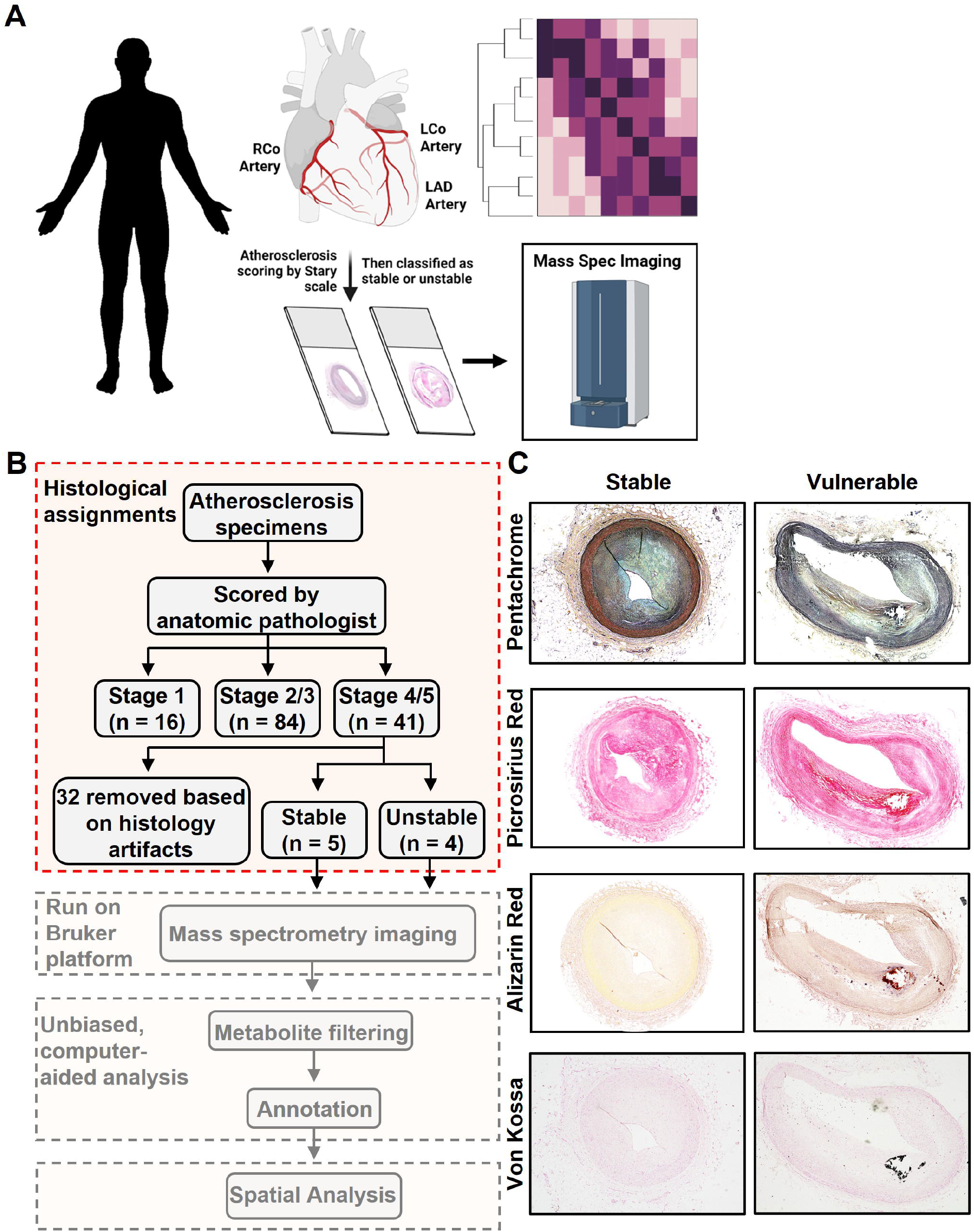
Experimental Design and Strategy of Mass Spectrometry Imaging of Stable and Unstable, Advanced Stage Human Atheromas. (A-B) We obtained 141 human atheromas and identified nine stage IV-V human atherosclerotic plaques suitable for MSI. Of these, four were determined to be unstable atheromas and five were stable atheromas. (C) Plaque staging and scoring for plaque vulnerability were aided by MOVAT Pentachrome, Picrosirius Red, Alizarin Red, and von Kossa staining. Representative images are shown for stable and unstable atheromas.

### Tissue histology

#### Von Kossa staining

Sections were deparaffinized in three changes of xylene and rehydrated through graded ethanol. To visualize calcium mineral deposition, rehydrated sections were incubated in 5% silver nitrate (Alfa Aesar, Haverhill, MA) under UV light for 20 minutes and then washed in distilled water. Sections were then washed with 5% sodium thiosulfate (MilliporeSigma, St. Louis, MO) and then distilled water to remove unreacted silver nitrate. Tissue specimens were counter-stained with 0.1% nuclear fast red (Acros Organics, Geel, Belgium), dehydrated in graded ethanol, cleared in xylene, then mounted using Permount™ Mounting Medium (Fisher Scientific, Hampton, NH), and imaged on a Keyence BZ-X800 (Keyence, Osaka, Japan).

#### Alizarin Red staining

To visualize deposited calcium, rehydrated sections were stained with 2% alizarin red S (Acros Organics, Geel, Belgium) for 5 minutes, followed by washing in 100% acetone, then acetone : xylene (1:1), cleared in 100% xylene, then mounted using Permount™ Mounting Medium (Fisher Scientific, Hampton, NH) and imaged on a Keyence BZ-X800 (Keyence, Osaka, Japan).

#### Picrosirius Red staining

Sections were deparaffinized in three changes of xylene and rehydrated through graded ethanol. Slides were placed in phosphomolybdic acid (Polysciences) for two minutes. After being rinsed in deionized water, sections were then placed in Picrosirius Red F3BA stain (Polysciences) for 45 minutes, followed by being incubated in 0.1N HCl for two minutes. Sections were placed in 70% ethanol, dehydrated, and mounted with a coverslip using Permount™ Mounting Medium (Fisher Scientific, Hampton, NH). Tissues were imaged on a Keyence BZ-X800 (Keyence, Osaka, Japan).

#### Pentachrome staining

To stain connective tissues, rehydrated sections were stained with a commercial Russel-Movat Pentachrome staining kit (Newcomer Supply, Middleton, WI), used following the manufacturer’s instructions. All sections were mounted using Permount™ Mounting Medium (Fisher Scientific, Hampton, NH) and imaged on a Keyence BZ-X800 (Keyence, Osaka, Japan).

### RNA-sequencing

RNA-sequencing (RNAseq) data in stable and unstable plaques from eight individuals with atherosclerosis were accessed from European Nucleotide Archive (PRJNA493259)^13^. Transcripts were quantified using Salman software package and consolidated at the gene level^14^. Differentially expressed genes (adjusted p < 0.05) between stable and unstable plaques were calculated with DESeq2^15^ and used in a gene set enrichment analysis for Gene Ontology (GO) biological processes with clusterProfiler^16^. Significantly enriched metabolic pathways (p < 0.05) were matched with MSI-identified compounds.

### Mass Spectrometry Imaging

Human artery cross-sections were deparaffinized in xylene, 2 × 3 min, and allowed to dry. Matrix was applied using an HTX M5 Robotic Reagent Sprayer (HTX Technologies, Chapel Hill, NC). For positive ion mode, 10 mg/mL α-cyano-4-hydroxycinnamic acid (CHCA) (Sigma-Aldrich, St. Louis, MO) in 50% acetonitrile, 0.1% trifluoroacetic acid was applied in 4 passes with a nozzle temperature of 75°C, a nozzle height of 40 mm, a flow rate of 120 µL/min, a track speed of 1200 mm/min, a track spacing of 3 mm, and a nitrogen pressure of 10 psi. For negative ion mode, 10 mg/mL 1,5-diaminonaphthalene (TCI America, Portland, OR) in 50% acetonitrile was applied in 10 passes with a nozzle temperature of 60°C, a nozzle height of 40 mm, a flow rate of 80 µL/min, a track speed of 1000 mm/min, a track spacing of 2 mm, and a nitrogen pressure of 10 psi. Images were acquired using Bruker FlexImaging 5.1 on a Bruker timsTOF fleX mass spectrometer (Bruker Daltonics, Billerica, MA) in both positive and negative ion mode at a spatial resolution of 30 µm over the *m/z* range 50-1000 with a Funnel 1 RF of 50 Vpp, a Funnel 2 RF of 100 Vpp, a Multipole RF of 150 Vpp, a Collision Energy of 3 eV, a Collision RF of 500 Vpp, a Transfer Time of 35 µs and a PrePulse Storage Time of 2 µs. A total of 1200 laser shots were summed per pixel. After data acquisition, the matrix was removed, and sections were stained with hematoxylin and eosin. Digital images of the sections were acquired at 20X magnification using a Hamamatsu NanoZoomerSQ Slide Scanner. Images were reviewed and annotated for areas of collagen-rich fibrous cap and necrotic core proximal area using NPD.view2 (Hamamatsu Corporation, Bridgewater, NJ). The annotated images were merged with images of the tissue sections used for data acquisition and regions of interest (ROIs) defined corresponding to the annotated regions. Images of all samples were combined and viewed in SCiLS Lab Pro 2021b (Bruker) after root mean squared normalization. Peaks were manually picked from the average spectrum for visualization and statistical analysis. T-tests and receiver operating characteristic analyses were carried out on the ROIs, comparing stable plaques to unstable plaques in the fibrous cap and necrotic core regions separately.

Putative metabolite identification was performed by loading the ROIs into MetaboScape 2021b (Bruker) and searching against the Human Metabolome Database (HMDB). Additionally, a smart formula search of the data was performed to determine potential chemical formulas for additional metabolites. Furthermore, *m/z* values were compared to metabolite image identifications in METASPACE (https://metaspace2020.eu/) or formulas queried against the Human Metabolome Database (https://hmdb.ca/). To integrate the MSI results with RNAseq of human stable and unstable atheromas (PRJNA493259)^13^, significantly enriched metabolic pathways identified through pathway enrichment analysis of RNAseq data were matched with MSI-identified compounds assigned to these pathways based on their metabolic identification and ontology described in HMDB.

### Metabolite Verification

To verify the identification of metabolites obtained from MetaboScape, METASPACE, and HMDB, signals observed in tissue sections were compared to purchased standards of target metabolites. Spectra of the standards were collected on both the Bruker timsTOF fleX and a Thermo Fusion Lumos Orbitrap (Thermo Corporation, San Jose, CA) mass spectrometer equipped with a MassTech AP/MALDI (ng) UHR source (MassTech, Inc., Columbia, MD). Accurate masses of the standards were compared to the signals observed in the imaging experiments. Furthermore, MS/MS was performed on the timsTOF fleX to determine characteristic fragments of the metabolite standards. The fragmentation patterns were compared to MS/MS spectra obtained directly from tissue sections. Identifications were verified based on accurate mass (<1 ppm) and matching fragmentation patterns.

### Statistical analysis

Statistical analyses were performed using GraphPad Prism version 9.0. Data were tested for normality using the Shapiro-Wilk test. If passed, Student’s *t*-test was used. Otherwise, Mann-Whitney U test was used. P < 0.05 was considered statistically significant.

## RESULTS

To select stable and unstable human atherosclerosis samples for mass spectrometry imaging (MSI) analysis, we utilized a biobank of atherosclerotic tissue specimens taken from the right coronary artery, left coronary artery, left anterior descending artery, and the carotid sinuses of 141 individuals. These tissues were formalin-fixed, paraffin-embedded, then sectioned and scored based on the Stary Classification scale^12^, then subdivided into stable and unstable atheromas. These samples were then processed for mass spectrometry imaging (MSI) and integrated with a publicly available RNAseq dataset comparing stable and unstable human atherosclerosis^13^. A general overview of the experimental design is described in **Figure 1A**, with the workflow of the MSI shown in **Figure 1B**. To score the atherosclerotic plaques, Pentachrome MOVAT staining, Picrosirius Red, Alizarin Red, and von Kossa staining were performed (**Figure 1C**). Based on the features from these stains, we identified 41 late-stage (stage 4/5) human atheromas. Because MSI demands high-quality tissue sections, we next removed samples that contained histological artifacts, such as wrinkles, tears, folds, or incomplete sectioning (n = 32). Based on fibrous cap thickness, we obtained five stable (fibrous cap > 65 microns) and four unstable (fibrous cap < 65 microns) atheromas (**Figure 1B**).

Cross-sections of these stable and unstable plaques were then deparaffinized, and matrix was applied using an automated reagent sprayer. Samples were then analyzed in positive and negative ion modes, and images were acquired using Bruker FlexImaging 5.1 on a Bruker timsTOF fleX mass spectrometer (**Figure 2A**) with a total of 1200 laser shots per pixel. Images of all samples were combined and viewed in SCiLS Lab Pro 2021b after root mean squared normalization. Two MSI examples of our collected human atherosclerosis specimens (thymine and homo-L-arginine) are shown in **Figure 2B**. The MALDI-MSI platform was able to identify 855 metabolites with assigned chemical formulas. We found that 77 metabolites were significantly elevated in the whole vessel of the stable plaques, and 179 metabolites were significantly elevated in the entire vessel of unstable plaques (**Figure 2C**). Metabolite identification was performed by loading the ROIs into MetaboScape 2021b (Bruker) and searching against software-deposited metabolite databases. Mass-to-charge values were also compared to metabolite image identifications in METASPACE (https://metaspace2020.eu/) followed by formula searching using the Human Metabolome Database (HMDB, https://hmdb.ca/) then queried for the designation of being previously detected and quantified (**Figure 3A**). This approach resulted in 189 annotated metabolites. To determine the rigor of our annotation strategy, we then selected twelve metabolites within different classes of molecules (e.g., nucleotides, amino acids, and fatty acids) and compared these to commercially purchased standards. Each standard was spotted to a MALDI target, and an MS/MS spectrum was collected using collision-induced dissociation (CID). The same parent *m/z* values observed in the standards were targeted in tissue sections and subjected to MS/MS. The fragments observed in the MS/MS spectra from tissues were compared to those obtained from the standards (**Supplemental Fig 1-4**, red arrows in each set of spectra indicate fragment peaks common between the standards and tissue). Unsurprisingly, it was found that more than one compound was fragmented simultaneously from tissue sections due to multiple peaks that were present within a ±1 Da isolation window, resulting in chimeric spectra with additional fragment peaks that do not entirely match the MS/MS spectra obtained from the standards. While several annotated xenobiotics could not be matched to standards (valproic acid, naproxen, salicylic acid), we were able to confidently assign all other tested metabolites, justifying the removal of xenobiotics from the list. This filtering system resulted in 170 annotated metabolites (**Figure 3B**).

**Figure 2.**
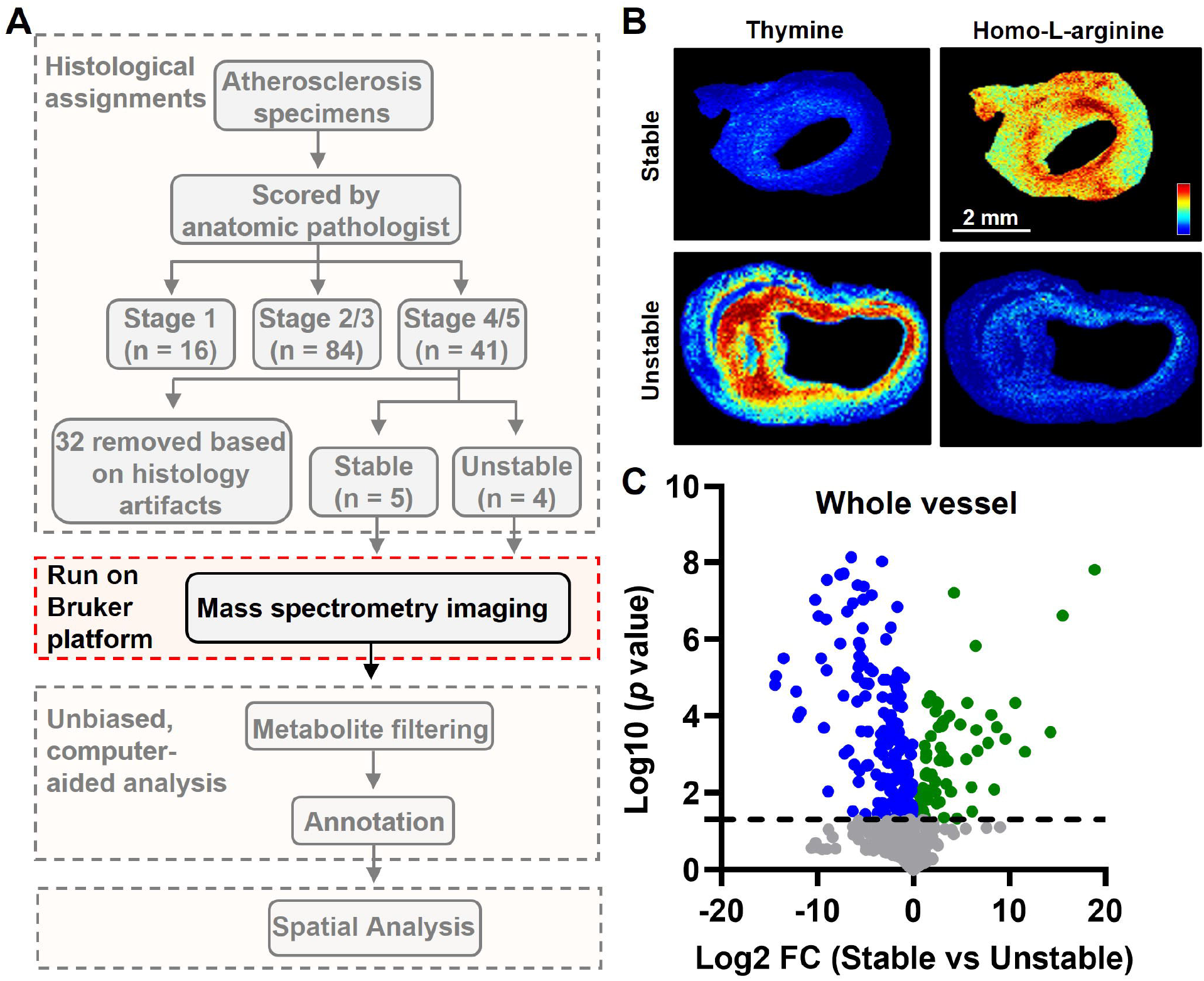
Mass Spectrometry Imaging of Human Atheromas. (A) MSI was performed after staging and scoring for plaque instability. (B) Matrix was applied to tissue sections using a Robotic Reagent Sprayer. 10 mg/mL α-cyano-4-hydroxycinnamic acid (CHCA) in 50% acetonitrile, 0.1% trifluoroacetic acid was used for positive mode, whereas 10 mg/mL 1,5-diaminonaphthalene in 50% acetonitrile was applied for negative ion mode. Images were acquired using Bruker FlexImaging 5.1 on a Bruker timsTOF fleX mass spectrometer in both positive and negative ion mode at a spatial resolution of 30 µm over the *m/z* range 50-1000. (B) Example images for thymine and homo-L-arginine are shown. (C) Data were displayed as a Volcano plot and significance was determined as log10 *p* values above 1.3 with a log2 fold change ± 1.5. Metabolites that are significantly elevated in the whole vessel of unstable plaques are displayed as blue circles, whereas metabolites significantly elevated in the whole vessel of stable plaques are displayed as green circles.

**Figure 3.**
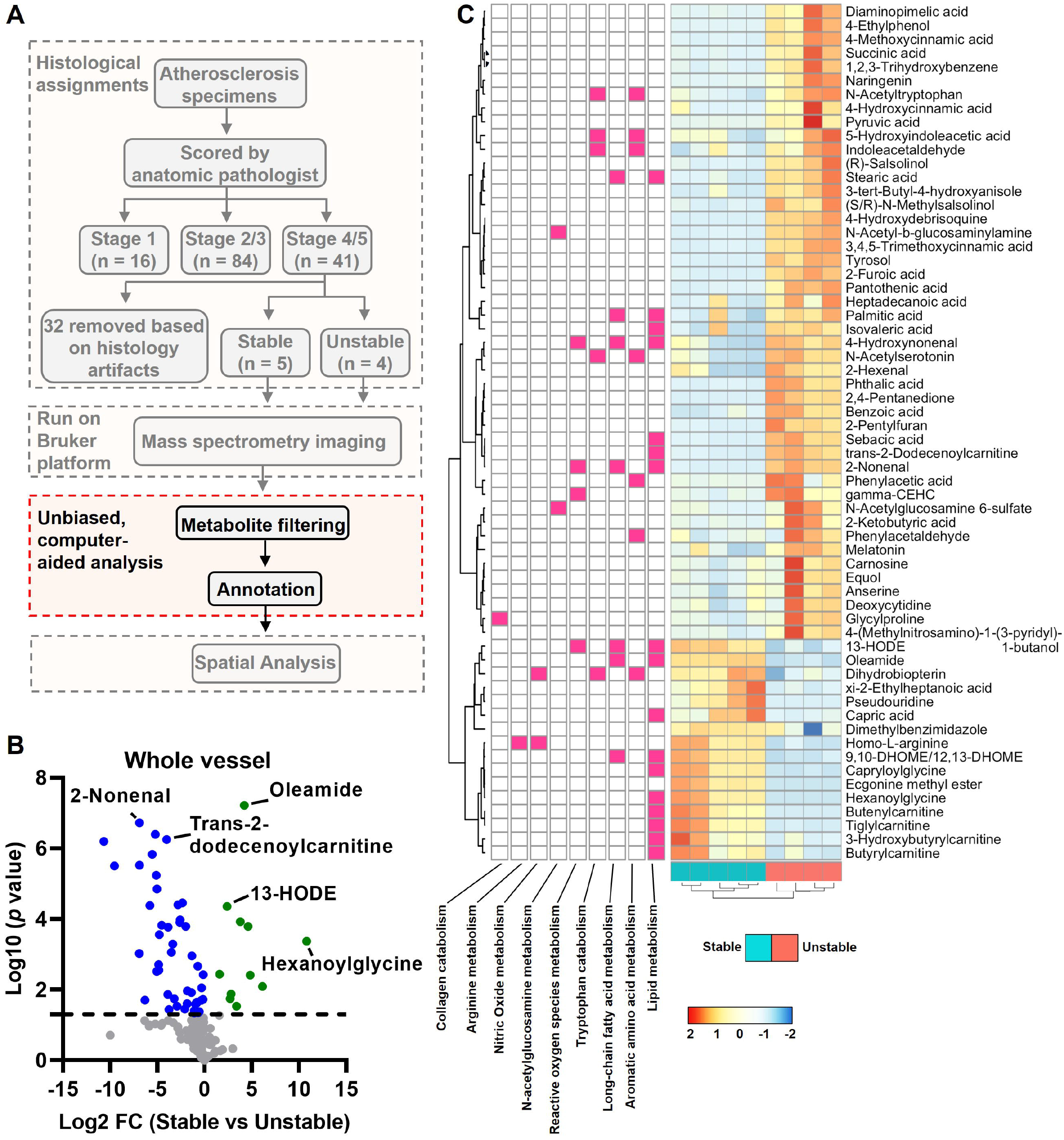
Metabolite Annotation and Integration with RNAseq of Stable and Unstable Atheromas. (A) Identification of metabolites was carried out by searching for deposited metabolites in MetaboScape. Mass-to-charge values were also compared to metabolite image identifications in METASPACE followed by formula searching using the Human Metabolome Database then queried for the designation of being detected and quantified. (B) Data from annotated metabolites are displayed as a Volcano plot and significance was determined as log10 *p* values above 1.3 with a log2 fold change ± 1.5. Metabolites that are significantly elevated in the whole vessel of unstable plaques are displayed as blue circles, whereas metabolites significantly elevated in the whole vessel of stable plaques are displayed as green circles. (C) Differential gene expression analysis was performed on an NCBI curated RNAseq dataset (GSE120521) comparing unstable and stable human atherosclerotic plaques. The impact of differentially expressed genes (*P* <0.05) on biological processes was determined by gene set enrichment analysis using the Gene Ontology database. Metabolic pathways that are significantly enriched (*P* < 0.05) are shown to the left of annotated metabolites.

We then coupled our annotated metabolites with RNAseq of human stable and unstable atheromas^13^, which revealed differences in pathways related to the metabolism of collagen, arginine, nitric oxide, N-acetylglucosamine, reactive oxygen species, aromatic amino acids, long-chain fatty acids, and lipids (**Figure 3C**). Twenty-nine of the annotated metabolites that were significantly different between stable and unstable atheromas could be assigned to at least one atherosclerosis-relevant pathway (**Figure 3C**). The pathway with the highest number of assigned metabolites was lipid metabolism. Whereas acylcarnitines (tiglylcarnitine, butenylcarnitine, butyrylcarnitine, and 3-hydroxybutyrylcarnitine) and acylglycines (hexanoylglycine and capryloylglycine), both known to be involved in fatty acid metabolism/oxidation, were enriched in stable plaques, saturated fatty acids (palmitic acid and stearic acid) and lipid peroxidation metabolites (4-hydroxynonenal and 2-nonenal) were increased in unstable plaques. Furthermore, aromatic amino acid and tryptophan metabolites (phenylacetic acid, phenylacetaldehyde, N-acetyltryptophan, N-acetylserotonin, indoleacetaldehyde, and 5-hydroxyindoleacetic acid) were predominantly found in unstable plaques. Notably, most metabolites could not be assigned to atherosclerosis-relevant pathways.

We next assessed differences in the fibrous cap and the necrotic core, comparing stable and unstable atheromas (**Figure 4A**). An example of two metabolites is shown (**Figure 4B**). In the fibrous cap, we identified 19 metabolites significantly elevated in unstable plaques, and 11 metabolites significantly elevated in stable plaques (**Figure 4C**). In the necrotic core microenvironment, we discovered 17 metabolites significantly elevated in the unstable atheromas and eight significantly elevated in the stable atheromas (**Figure 4D**). Interestingly, homo-L-arginine, a substrate for nitric oxide synthase and an emerging protective cardiovascular risk factor^17-19^, was enriched in the fibrous cap and necrotic core of stable atheromas (**Figure 4C and D**). Whereas the lipid peroxidation metabolite 2-nonenal was elevated in both the fibrous cap and necrotic core of unstable atheromas (**Figure 4C and D**), pantothenic acid (vitamin B5), which contributes to fatty acid synthesis and reduces oxidative stress^20,21^, was enriched only in the fibrous cap (**Figure 4C**). We next assessed metabolite differences in the fibrous cap versus the necrotic core. In stable plaques, pyruvic acid was significantly increased in the fibrous cap, whereas lactic acid was increased in the necrotic core (**Figure 4E**). In unstable plaques, the tryptophan metabolite 5-hydroxyindoleacetic acid, linked to metabolic syndrome and systemic inflammation^22^, was significantly increased in the fibrous cap (**Figure 4F**). Altogether, using MALDI-MSI coupled with transcriptomics, we identified metabolic differences in the whole vessel, fibrous cap, and necrotic core between stable and unstable atheromas.

**Figure 4.**
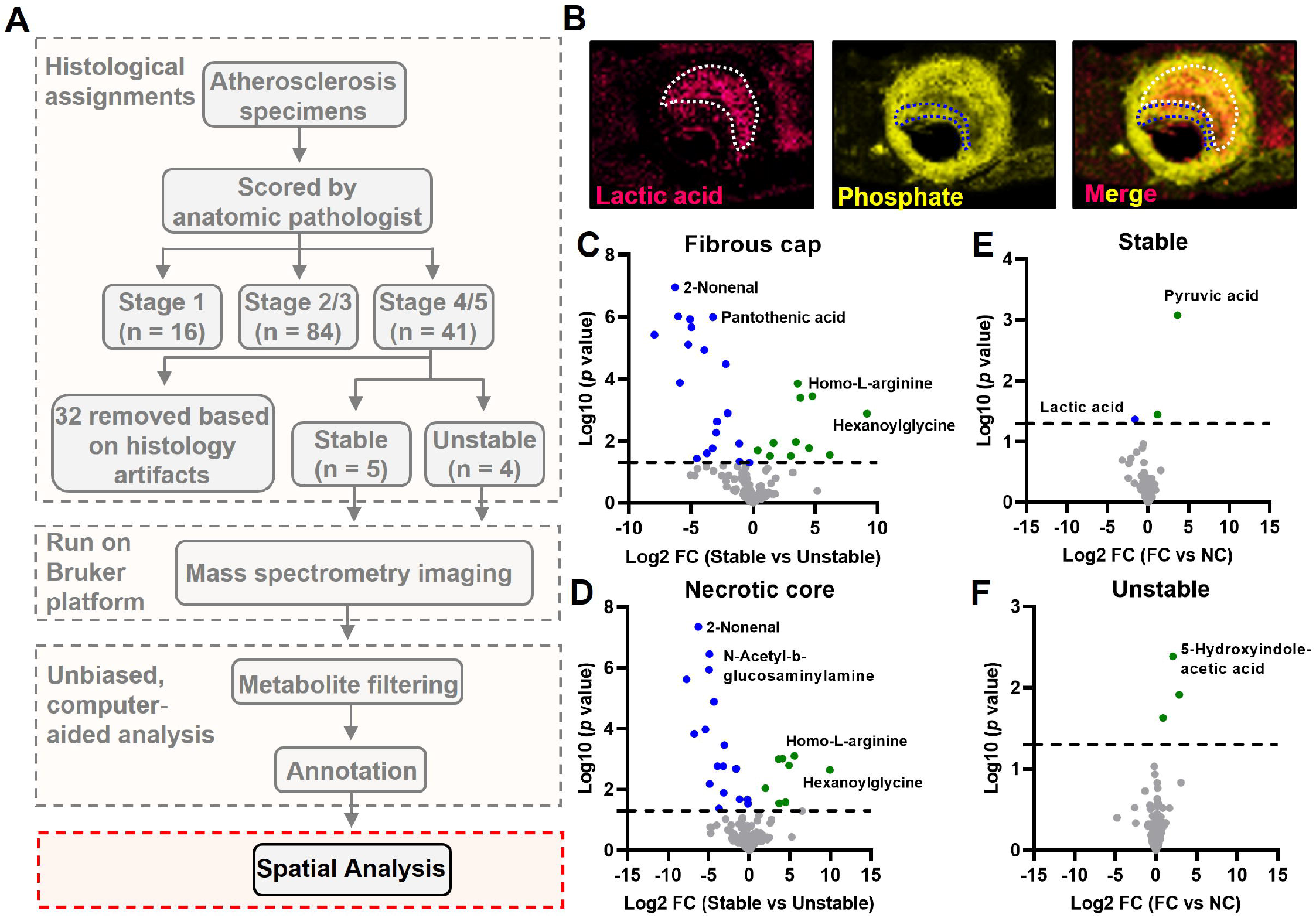
Spatial Resolution of Metabolites in the Fibrous Cap and Necrotic Core of Stable and Unstable Human Atheromas. (A) Spatial determination of annotated metabolites from Figure 3B was performed. (B) Images of lactic acid and phosphate are shown as examples of metabolites enriched in the fibrous cap and necrotic core. Data from annotated metabolites are displayed as a Volcano plot and significance was determined as log10 *p* values above 1.3 with a log2 fold change ± 1.5. Metabolites that are significantly elevated in the whole vessel of unstable plaques are displayed as blue circles, whereas metabolites significantly elevated in the whole vessel of stable plaques are displayed as green circles. (C-F) Metabolites that are significantly elevated in the fibrous cap (C) and necrotic core (D) of unstable plaques are displayed as blue circles, whereas metabolites significantly elevated in stable plaques are displayed as green circles. In stable plaques (E) and unstable plaques (F), metabolites that are significantly elevated in the fibrous cap are displayed as green circles, whereas metabolites significantly elevated in the necrotic core are displayed as blue circles. Data from metabolites are displayed as a Volcano plot and significance was determined as log10 *p* values above 1.3 with a log2 fold change ± 1.5.

## DISCUSSION

Elegant omics studies, including RNAseq, scRNAseq, and bulk metabolomics, have revealed differences in gene expression, cell composition, and metabolite profiles in human atherosclerosis^23,24^. However, these studies do not inform us wherein the plaques these differences occur. Here, we aimed to spatially resolve metabolites within the necrotic cores and fibrous caps of stable and unstable human atheromas in an unbiased manner. We annotated 170 out of 855 metabolites and integrated these results with a publicly available RNAseq dataset that determined a majority of annotated metabolites belonged to metabolic pathways that are not currently associated with atherosclerosis. We also found a variety of classes of metabolites in the fibrous cap and necrotic cores of stable and unstable atheromas.

Our dataset revealed that acylcarnitines and acylglycines were enriched in stable plaques. While our data might suggest that enhancing acylcarnitine levels would be atheroprotective, L-carnitine supplements, which enhance acylcarnitine levels^25^, worsen stenosis in individuals with metabolic syndrome^26^. Furthermore, links between elevated acylcarnitines and insulin resistance have been described^27,28^. Also, elevations in acylcarnitines are associated with an increased risk of acute cardiovascular events^29-31^. In contrast to acylcarnitines, increased circulating glycine levels are linked to protections in clinical and experimental atherosclerosis^9,32,33^, and enhanced glycine levels increase circulating levels of acylglycines^34^. These fatty acid-conjugated amino acids, particularly acylcarnitines, are critical for transporting fatty acids across the inner mitochondrial membrane^35^, and the presence of these fatty acid-conjugated amino acids in stable plaques is in line with enhancements in fatty acid oxidation, an atheroprotective metabolic pathway^4,5^.

In contrast to acylcarnitines and acylglycines, saturated fatty acids (palmitic acid and stearic acid) were found to be increased in unstable plaques. Data from the Rotterdam study revealed that intake of palmitic acid was strongly associated with coronary artery disease^36^. Consistently, feeding mice a palmitic acid-enriched diet modulates vascular smooth muscle cell (vSMC) phenotype^37^, drives inflammasome activation^38^, and enhances reactive oxygen species^39^, cellular processes that drive vascular calcification and plaque instability^40,41^. In contrast to palmitic acid, the role of stearic acid in atherosclerosis remains inconclusive. For instance, stearic acid not only lowers LDL-cholesterol but also lowers HDL-cholesterol^42^. Also, dietary stearic acid does not increase atherosclerosis risk and is linked with reductions in systolic blood pressure and improvements in cardiac function^43-45^. However, stearic acid accumulation in macrophages induces endoplasmic reticulum stress-mediated apoptosis^46^, a process known to drive necrotic core formation and plaque instability. While stearic acid alone may not impact one’s risk of atherosclerosis, the action of both stearic acid and palmitic acid may drive vSMC phenotypic modulation, enhance proinflammatory responses in macrophages, and increase apoptosis.

In addition to saturated fatty acids, aromatic amino acids and tryptophan metabolites were also enriched in unstable plaques. Interestingly, phenylacetic acid induces reactive oxygen species in vSMCs and TNFα production in endothelial cells^47,48^. The fatty acid-conjugated amino acid N-acetyltryptophan is modified from tryptophan by the gut microbiota that is then absorbed by the intestine^49^. Supporting our observation that N-acetyltryptophan was present in unstable atheromas, enhanced levels of N-acetyltryptophan are associated with an increased risk of coronary heart disease^50^, which may be attributed to the effect of N-acetyltryptophan on hyperlipidemia^51^. We found that the serotonin derivative 5-hydroxyindoleacetic acid (5-HIAA) was enriched in the fibrous cap of unstable plaques, and interestingly, 5-HIAA is associated with metabolic syndrome and C-reactive protein (CRP), a biomarker of inflammation^22^. 5-HIAA can be oxidized by myeloperoxidase to form reactive quinones, and these 5-HIAA reactive quinones are found in advanced human atherosclerotic lesions^52^. However, the role of 5-HIAA in atherosclerosis has yet to be explored.

An exciting finding revealed by our studies was the distinct locations of pyruvic acid and lactic acid in the fibrous cap and necrotic core of stable plaques, respectively. During acute and chronic inflammation, a shift in immune cell metabolism towards glycolysis that leads extracellular lactate accumulation occurs^53^, and elevations in circulating lactate are associated with hypertension^54^, type 2 diabetes^55^, and carotid atherosclerosis^56^. Our findings that revealed lactate is enriched in the necrotic core support this concept that macrophages localized in these regions are proinflammatory. However, instead of promoting inflammatory responses that drive atherosclerosis progression, lactate plays a crucial role in resolution by driving IL-10 production in phagocytes engulfing apoptotic cells^57^, inducing histone lactylation^58^, and suppressing proinflammatory responses by inhibiting YAP and NFκB activation in macrophages^59^. In contrast to lactate, we found pyruvate was enriched in the fibrous caps of stable plaques, and pyruvate has been linked to metabolic dysregulations and associated with worsening atherosclerosis. Consistently, pyruvate seems to promote features that drive fibrous cap formation. For instance, deleting pyruvate kinase M2, which substantially reduces the formation of pyruvate, in vSMCs suppresses phenotypic modulation and attenuates neointimal hyperplasia^60^.

Our MSI data presented here validate known metabolites associated with atherosclerosis progression while also pointing to classes of metabolites that may play significant roles in plaque instability. Importantly, we were unable to confidently annotate the majority of metabolite-related peaks that we identified. However, future technical innovations that permit accurate annotation of these peaks will undoubtedly reveal more metabolites that play critical roles in plaque instability. Thus, we view our work presented here as a first step to defining an atlas of metabolic pathways involved in plaque destabilization in human atherosclerosis. We anticipate that this work will be a valuable resource for the scientific community and will ultimately open new avenues of research in cardiovascular disease.

## ACKNOWLEDGEMENTS

We thank De’Lacy Lewis for her assistance with the immunofluorescence histochemistry staining and Arun Sreekumar for donation of metabolite standards.

