## Supplementary Figures 1-4 for "Spatially Resolved Metabolites in Stable and Vulnerable Human Atherosclerotic Plaques Identified by Mass Spectrometry Imaging"

### Supplemental Figure 1

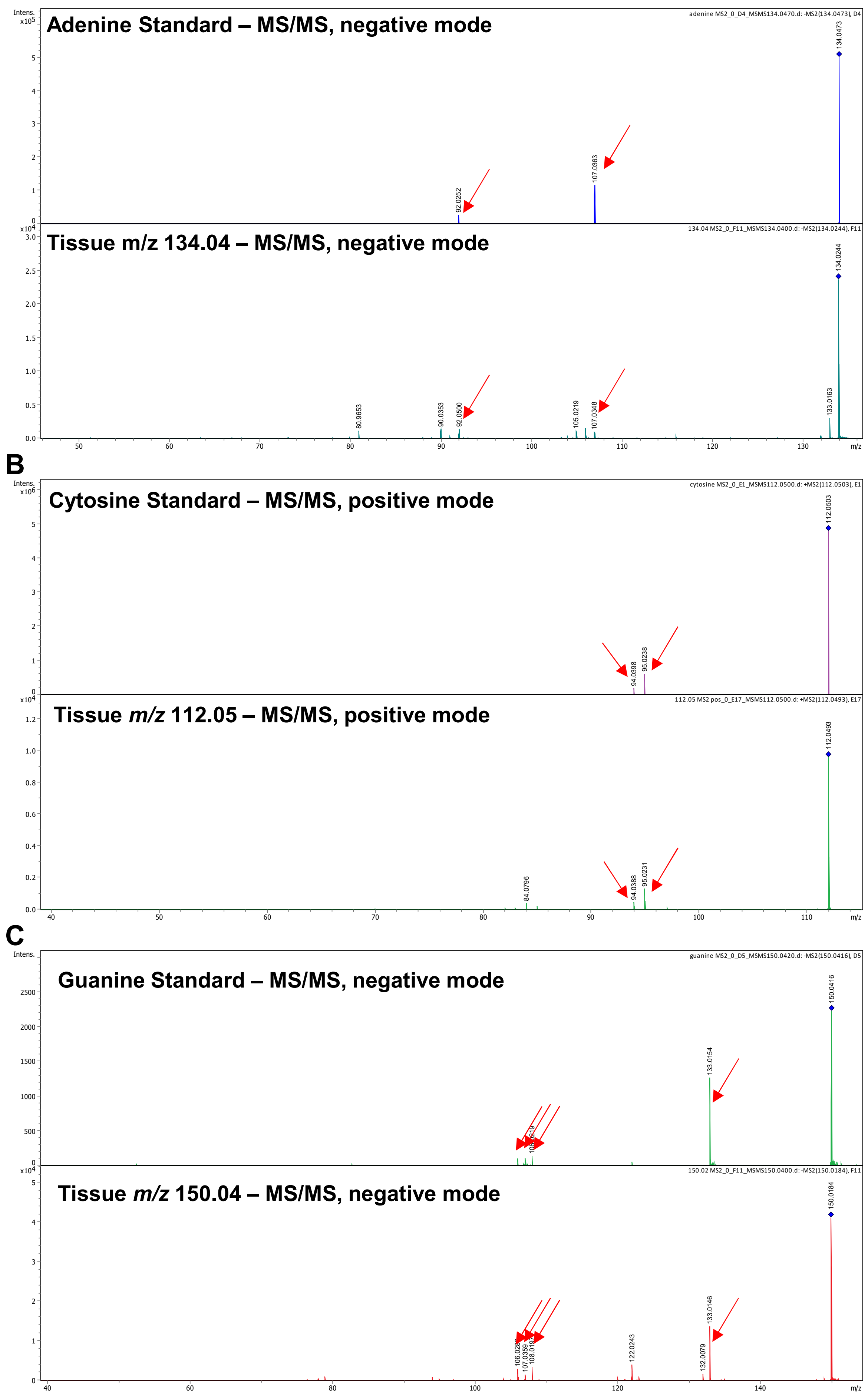

**Supplemental Figure 1. Confirmation of Nucleotides using Analytic Standards.** (**A**) Standards for adenine, (**B**) cytosine, and (**C**) guanine were spotted to a MALDI target and an MS/MS spectrum was collected using collision-induced dissociation (CID). Parent *m/z* values were observed in the standards were evaluated in the tissue sections. The fragments observed from the tissue were compared to those obtained from the standards. Red arrows in each spectra indicate fragment peaks common between standards and tissue.

### Supplemental Figure 2

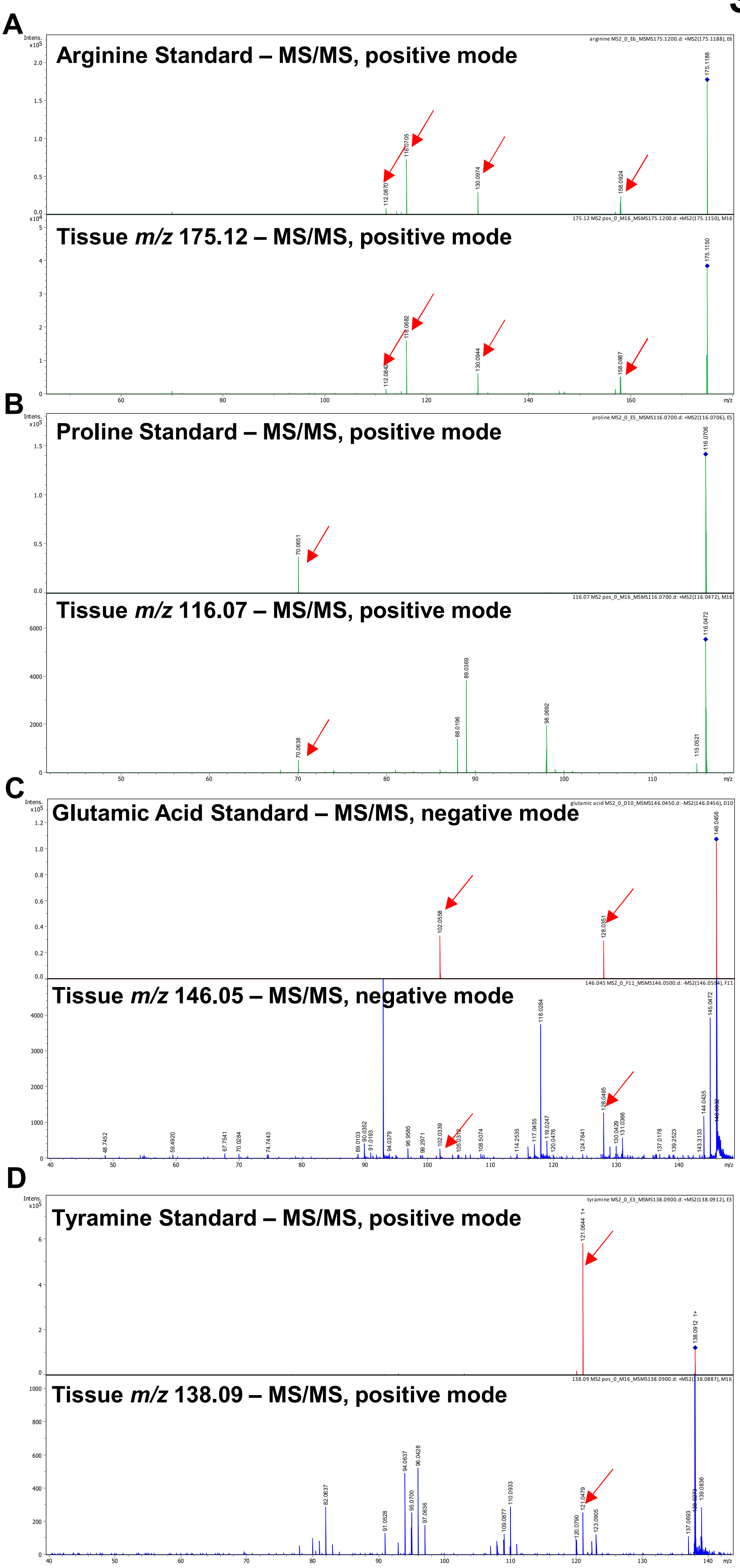

### Supplemental Figure 2
### Figure Legend

**Supplemental Figure 2. Confirmation of Amino Acids using Analytic Standards.** (**A**) Standards for arginine, (**B**) proline, (**C**) glutamic acid, (**D**) and tyramine were spotted to a MALDI target, and an MS/MS spectrum was collected using CID. Parent *m/z* values were observed in the standards were evaluated in the tissue sections. The fragments observed from the tissue were compared to those obtained from the standards. Red arrows in each spectra indicate fragment peaks common between standards and tissue.

### Supplemental Figure 3

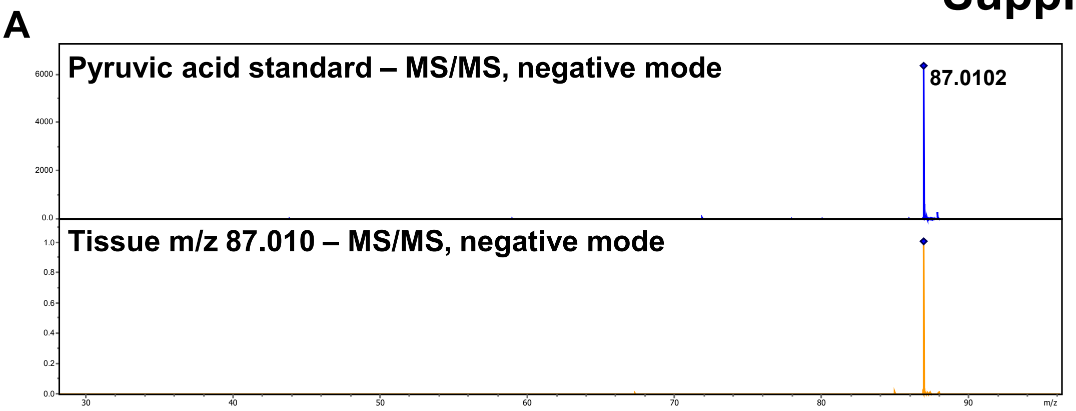


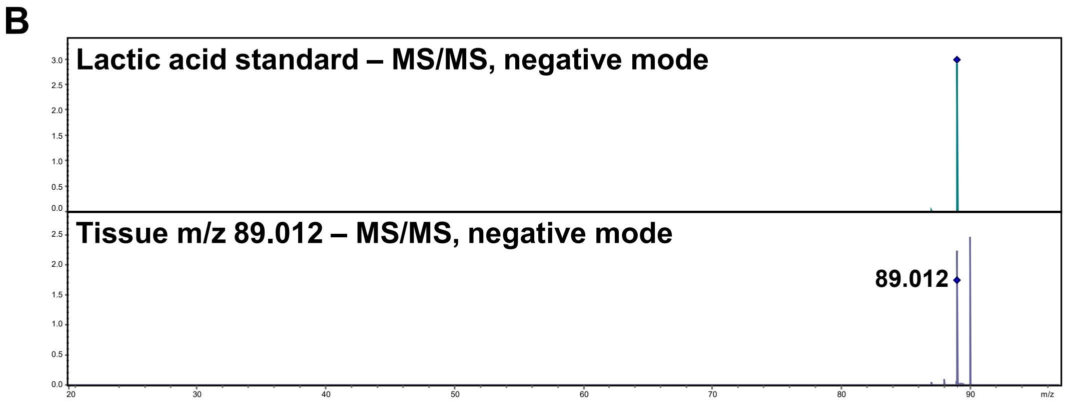

### Supplemental Figure 3
### Figure Legend

**Supplemental Figure 2. Confirmation of Pyruvic Acid and Lactic Acid using Analytic Standards.** (**A**) Standards for pyruvic acid and (**B**) lactic acid were spotted to a MALDI target, and an MS/MS spectrum was collected using CID. Parent *m/z* values were observed in the standards were evaluated in the tissue sections. The fragments observed from the tissue were compared to those obtained from the standards.

### Supplemental Figure 4

A

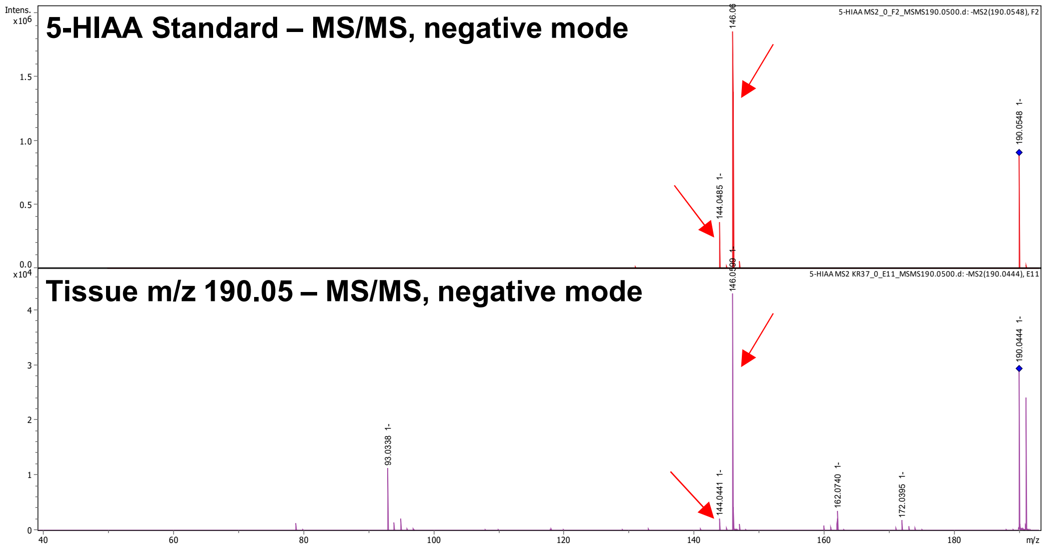


B

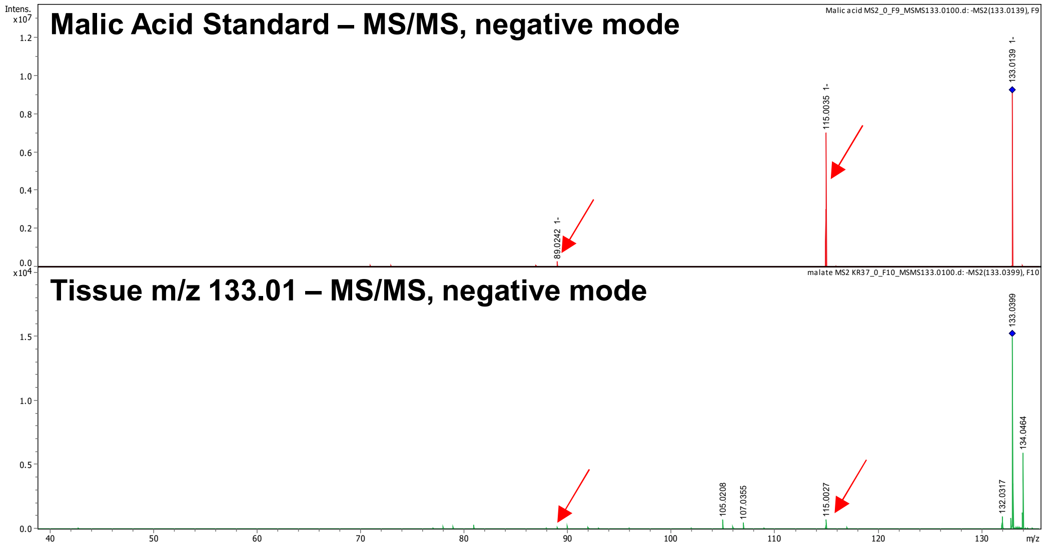


C

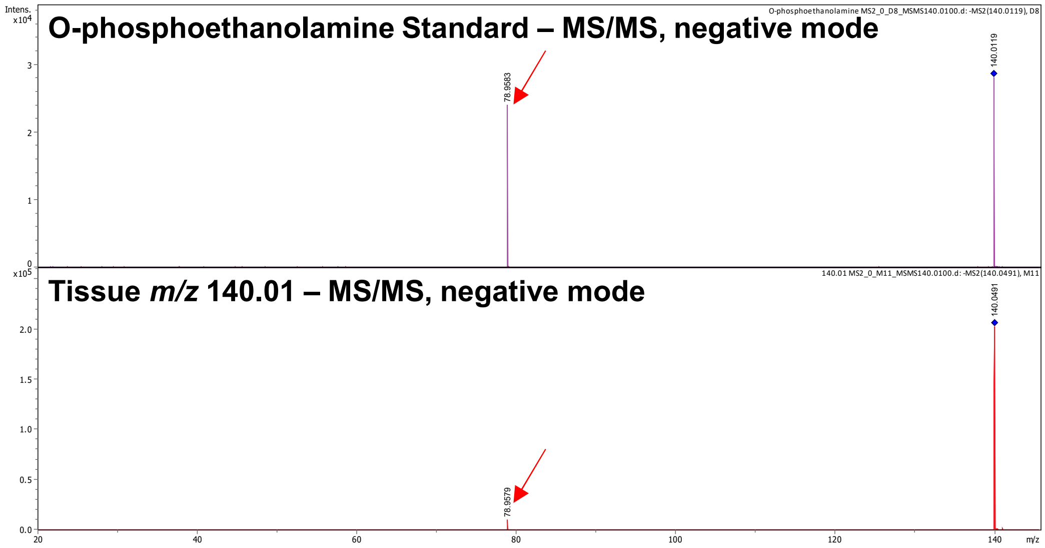

### Supplemental Figure 4
### Figure Legend

**Supplemental Figure 4. Confirmation of 5-HIAA, Malic Acid, and O-Phosphoethanolamine using Analytic Standards.** (**A**) Standards for 5-HIAA, (**B**) malic acid, (**C**) O-phosphoethanolamine were spotted to a MALDI target, and an MS/MS spectrum was collected using CID. Parent *m/z* values were observed in the standards were evaluated in the tissue sections. The fragments observed from the tissue were compared to those obtained from the standards. Red arrows in each spectra indicate fragment peaks common between standards and tissue.
